# VAMP8 contributes to TRIM6-mediated type-I interferon antiviral response during West Nile virus infection

**DOI:** 10.1101/749853

**Authors:** Sarah van Tol, Colm Atkins, Preeti Bharaj, Kendra N. Johnson, Adam Hage, Alexander N. Freiberg, Ricardo Rajsbaum

**Author notes:** Address correspondence to: Ricardo Rajsbaum, Alexander N. Freiberg. C.A and S.v.T. contributed equally to this work.

## Abstract

Several members of the tripartite motif (TRIM) family of E3 ubiquitin ligases regulate immune pathways including the antiviral type I interferon (IFN-I) system. Previously, we demonstrated that TRIM6 is involved in IFN-I induction and signaling. In the absence of TRIM6, optimal IFN-I signaling is reduced, allowing increased replication of interferon-sensitive viruses. Despite having evolved numerous mechanisms to restrict the vertebrate host’s IFN-I response, West Nile Virus (WNV) replication is sensitive to pre-treatment with IFN-I. However, the regulators and products of the IFN-I pathway that are important in regulating WNV replication are incompletely defined. Consistent with WNV’s sensitivity to IFN-I, we found that in TRIM6 knockout (TRIM6-KO) A549 cells WNV replication is significantly increased and IFN-I induction and signaling is impaired compared to wild-type (wt) cells. IFNβ pre-treatment was more effective in protecting against subsequent WNV infection in wt cells as compared to TRIM6-KO, indicating that TRIM6 contributes to the establishment of an IFN-induced antiviral response against WNV. Using next generation sequencing, we identified VAMP8 as a potential factor involved in this TRIM6-mediated antiviral response. VAMP8 knockdown resulted in reduced Jak1 and STAT1 phosphorylation and impaired induction of several ISGs following WNV infection or IFNβ treatment. Furthermore, VAMP8-mediated STAT1 phosphorylation required the presence of TRIM6. Therefore, the VAMP8 protein is a novel regulator of IFN-I signaling, and its expression and function is dependent on TRIM6 activity. Overall, these results provide evidence that TRIM6 contributes to the antiviral response against WNV and identified VAMP8 as a novel regulator of the IFN-I system.

**IMPORTANCE:** WNV is a mosquito-borne flavivirus that poses threat to human health across large discontinuous areas throughout the world. Infection with WNV results in febrile illness, which can progress to severe neurological disease. Currently, there are no approved treatment options to control WNV infection. Understanding the cellular immune responses that regulate viral replication is important in diversifying the resources available to control WNV. Here we show that the elimination of TRIM6 in human cells results in an increase in WNV replication and alters the expression and function of other components of the IFN-I pathway through VAMP8. Dissecting the interactions between WNV and host defenses both informs basic molecular virology and promotes the development of host- and viral-targeted antiviral strategies.

## INTRODUCTION

West Nile Virus (WNV) is an enveloped positive sense single stranded RNA virus and a member of the family *Flaviviridae* (1, 2). Mosquitoes competent for WNV (predominantly *Culex*) transmit the virus through blood feeding (3). Enzootic transmission cycles between birds and mosquitoes maintain the virus in the environment, but mosquitoes also incidentally infect humans and other mammals that act as dead-end hosts. In 1999, WNV was introduced to North America and has since then become an endemic pathogen, causing annual outbreaks in human populations, and is the leading cause of mosquito-borne encephalitis (4). Although primarily asymptomatic, WNV infection causes flu-like symptoms in approximately 20% of infected humans with a fewer than 1% of symptomatic cases progressing to neurologic manifestation (5). The case fatality rate for symptomatic cases is approximately 10% (1). Currently, no WNV vaccines or anti-viral treatments are approved for human use (6–10).

Understanding the molecular mechanisms of WNV replication at the host cellular level, and specifically WNV-host type I interferon (IFN-I) interactions, may allow identifying targets for antiviral development. Many groups have demonstrated that interferon-stimulated gene (ISG) products, such as ISG54 (IFIT2) (11), IFITM3 (12), and Oas1b (13), and others (Reviewed in: (14)) restrict WNV replication. Further, in mouse models of WNV infection, lack of IFN-I induction through signaling factors such as TLR3 (15), IRF7 (16), RIG-I (17), IFNβ (18), IFNAR (16), STAT1 (19), and IKKε (11) increases susceptibility to WNV. Mutations in IFN-I pathway genes and ISGs have also been associated with increased disease during WNV infections in humans (20, 21). Despite WNV’s sensitivity to IFN-I, WNV has evolved several mechanisms to antagonize IFN-I including NS1 interference with RIG-I and MDA5 function (22), NS3 helicase impairment of Oas1b activity (13), and NS5 disruption of the type-I interferon receptor (IFNAR) surface expression (23), STAT1 phosphorylation (24), and sub-genomic flavivirus RNA (25). Since WNV’s resistance to IFN-I contributes to virulence (16, 26), defining the IFN-I signaling pathway components required to respond to WNV infection is important in aiding the development of WNV-specific therapies.

Upon WNV infection, pathogen recognition receptors including RIG-I and MDA5 recognize viral RNA (17) and signal through their adaptor MAVS to activate downstream IKK-like kinases TBK1 and IKKε. Activation of TBK1 and IKKε promote IFN-I production through activation of transcription factors IRF3 and IRF7 (27). IFN-I is then secreted and engages the IFN-I receptor to induce IFN-I signaling. Early in the IFN-I signaling cascade, the kinases Jak1 and Tyk2 phosphorylate STAT1 (Y701) and STAT2 (Y690), which dimerize and interact with IRF9 to form the ISGF3 complex (28, 29). ISGF3 then translocates to the nucleus where it interacts with interferon-stimulated response element (ISRE) present in the promoter of ISGs. In addition, upon IFN-I stimulation, IKKε plays an essential non-redundant role in phosphorylation of STAT1 on S708, which is required for induction of IKKε-dependent ISGs (30). Several IKKε-dependent ISGs, including ISG54 (30), are involved in restricting WNV (11).

TRIM6, an E3 ubiquitin ligase in the tripartite motif (TRIM) protein family, plays a crucial role in facilitating the activation of the IKKε-dependent branch of the IFN-I signaling pathway (31). In concert with the ubiquitin activating (E1) enzyme and the ubiquitin conjugating (E2) enzyme, UbE2K, TRIM6 synthesizes unanchored K48-linked polyubiquitin chains that promote the oligomerization and autophosphorylation of IKKε (T501) (31). Following phosphorylation of T501, IKKε is activated and phosphorylates STAT1 (S708) to promote the transcription of the IKKε-dependent ISGs (30, 31). Due to TRIM6’s role in activating IKKε-dependent IFN-I signaling and the importance of IKKε-specific ISGs in restricting WNV, we hypothesized that WNV replication would be enhanced in the absence of TRIM6.

Here we show that WNV viral load is increased and that IFN-I induction and IKKε-dependent IFN-I signaling are impaired in TRIM6-knockout (TRIM6-KO) cells. Next-generation RNA sequencing (NGS) identified several ISGs expressed at lower levels in TRIM6-KO cells compared to wild-type (wt) cells, including several ISGs previously described to restrict WNV replication. In addition to ISGs, the absence of TRIM6 also affects the expression of genes not known to regulate WNV replication of IFN-I signaling, including *Vamp8*. We investigated the role of a TRIM6-dependent gene not previously described to regulate WNV replication or IFN-I signaling, *Vamp8*. VAMP8 is a vesicle-associated-membrane protein in the SNARE (Soluble-*N*-ethylmaleimide-sensitive factor attachment protein) family known to modulate endocytosis (32), exocytosis of secretory (33–36) and lytic granules (37), thymus development (38), receptor exocytosis (39), and cross-presentation by antigen presenting cells (40). We found that VAMP8 does not directly affect WNV replication but promotes optimal IFN-I signaling. Overall, we conclude that TRIM6 is important in promoting optimal IFN-I induction and signaling during WNV infection and that VAMP8 is a novel TRIM6-dependent factor involved in regulating IFN-I signaling.

## RESULTS

### WNV Replication is Increased in IFN-I Induction and Signaling Impaired TRIM6-KO Cells

To test our hypothesis that the absence of TRIM6 facilitates WNV replication, growth kinetics at different multiplicity of infections (MOI) were determined in wt and TRIM6-KO A549 cells. A significant increase in viral replication was detected in TRIM6-KO cells at 48 hours post-infection (h.p.i.) in comparison to wt cells when infections were performed at both low (0.1 PFU/cell) and high (5.0 PFU/cell) MOIs (Figure 1A-1B). To address the effect of the absence of TRIM6 on the IFN-I pathway, protein expression and phosphorylation of IFN-I pathway components in WNV infected cells were assessed (Figure 1C). Activation of both TBK1 and IKKε kinases are known to contribute to efficient IRF3 phosphorylation and IFN-I induction. While the levels of phosphorylated TBK1 (S172), were substantially higher in TRIM6-KO than in wt cells at late time points p.i. (Figures 1C, S1A), the TRIM6-dependent phosphorylation on IKKε-T501 was significantly attenuated in TRIM6-KO cells (Figures 1C, S1C), as we previously reported in a model of Influenza infection (31). Although less striking, phosphorylation on IKKε-S172 was also reduced in TRIM6-KO cells (Figures 1C, S1B). These data suggest that TRIM6 is important for promoting efficient activation of IKKε but not TBK1 in the context of WNV infection.

**Figure 1.**
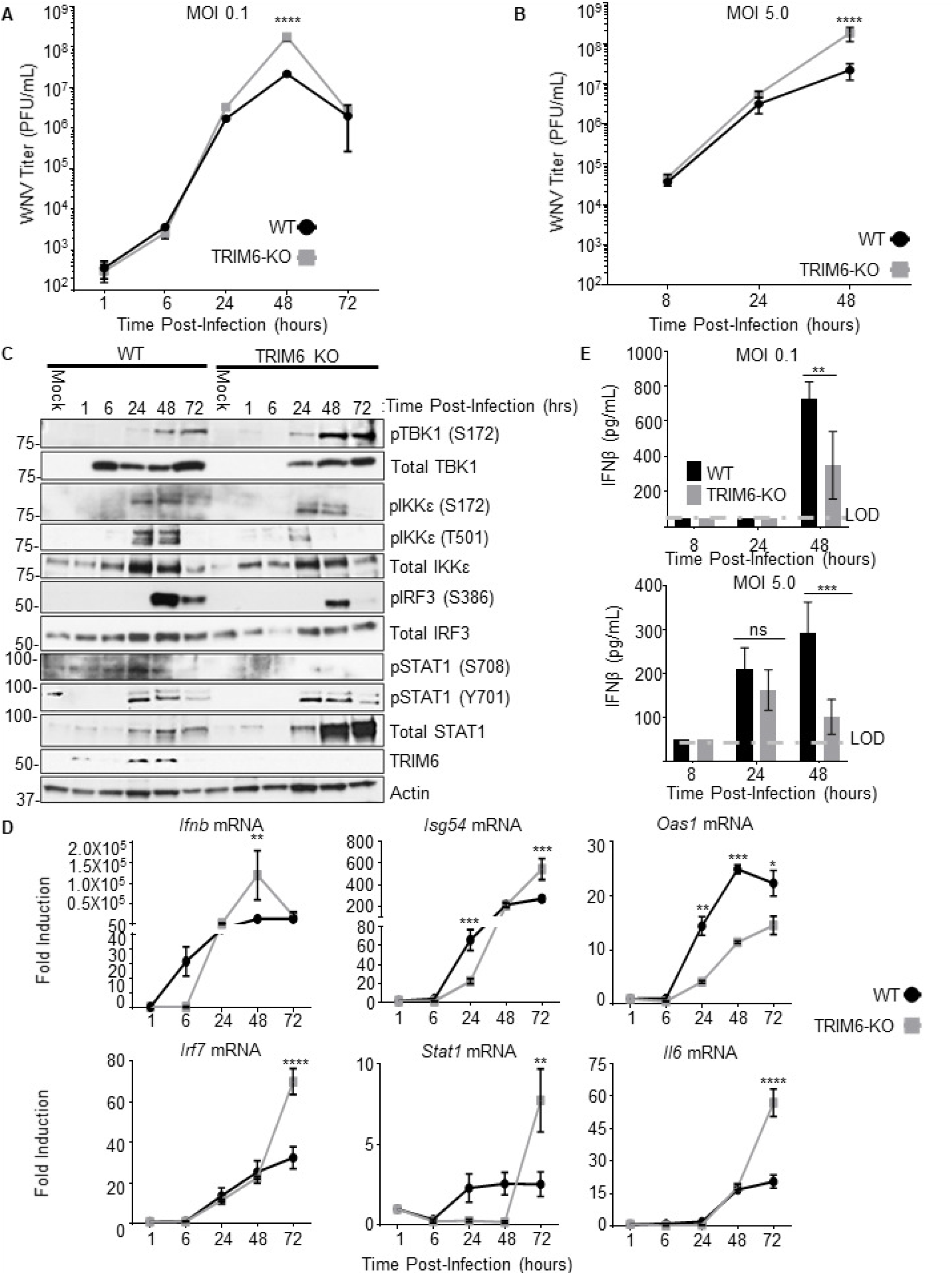
Increased WNV Replication in TRIM6-KO Cells is Associated with Impaired IFN-I Induction and Signaling. wt or TRIM6-KO A549s were infected with WNV 385-99 at MOI 0.1 (A, C, E top) or 5.0 (B, D, E bottom). The viral load in supernatants of infected cells was measured by plaque assay on Vero CCL-81 cells (A, B). Whole cell lysates from WNV (MOI 0.1) infected cells were run on western blot for analysis of protein expression and phosphorylation (C). RNA isolated from mock and WNV-infected cells was isolated to assess gene expression of *Ifnb*, ISGs: *Isg54*, *Oas1*, *Irf7 and Stat1*, and a non-IFN-I regulated gene: *Il6* (D). Change in expression represented as fold induction (D). IFNβ was measured via ELISA of irradiated supernatants infected with WNV at MOIs 0.1 and 5.0 (E). Error bars represent standard deviation (n=3). For statistical analysis, two-way ANOVA with Tukey’s post-test for multiple comparisons was used; ****p <0.0001, ***p <0.001, **p <0.01, *p <0.05. All experiments were performed in triplicate and immunoblots are representative sample. All experiments were repeated at least 2 times.

Although the amount of pTBK1 (S172) is increased in TRIM6-KO cells, the phosphorylation of IRF3 (S386) is lower in TRIM6-KO compared to wt cells (Figures 1C, S1D), suggesting that TBK1 can not completely compensate for reduced IKKε activity under these conditions of WNV infection. In line with impaired IRF3 activation in TRIM6-KO cells, *Ifnb* mRNA levels were reduced in TRIM6-KO cells only at early time points post-infection (6 hpi). However, the expression levels of *Ifnb* in TRIM6-KO cells, significantly increased over time as compared to WT cells and correlated with the increased virus titers at 48 h.p.i. (Figure 1D). Despite the increase in *Ifnb* mRNA levels at 48 hpi in TRIM6-KO cells, the amount of secreted IFNβ protein was significantly decreased in TRIM6-KO compared to wild-type cells (Figure 1E).

Consistent with reduced IFNβ protein accumulated in supernatants of infected TRIM6-KO cells, the amount of pSTAT1 (Y701) is lower in TRIM6-KO cells compared to wt at all time points evaluated (Figures 1C, S1E). In contrast, the expression of total STAT1 is substantially increased in TRIM6-KO cells compared to wt, which is consistent with the reported accumulation of unphosphorylated STAT1 in IKKε-KO cells (41). Phosphorylation on STAT1 (S708), an IKKε and TRIM6-dependent modification (30, 31), is nearly undetectable in the TRIM6-KO cells upon WNV infection (Figure 1C, S1F).

Consistent with this defect in the TRIM6-IKKε branch of the IFN-I signaling pathway in TRIM6-KO cells, the TRIM6/IKKε-dependent ISGs *Isg54* and *Oas1* (30, 31), have different patterns of induction. In the case of *Isg54*, induction is significantly lower in the TRIM6-KO cells than in wt cells at 24 hpi, but this pattern is reversed at 72 hpi mirroring *Ifnb* expression (Figure 1E). In contrast, induction of *Oas1* is attenuated in TRIM6-KO cells 24-72 hpi (Figure 1E). IKKε-independent ISGs *Irf7* and *Stat1* and non-ISG *Il-6* are expressed at higher levels in TRIM6-KO cells at later time points (Figure 1E), again in correlation with *Ifnb* induction. As pIKKε (T501) plays a non-redundant role in facilitating the expression of multiple ISGs (30, 31), that are known to be important in restricting WNV replication (11, 13, 42), the reduced capacity of TRIM6-KO cells to phosphorylate this residue likely contributes to the impaired antiviral activity against WNV. Overall, the absence of TRIM6 augments WNV replication and impairs the IKKε-dependent branch of the IFN-I pathway in line with our previous findings with other viruses including Influenza (IAV), Sendai (SeV) and Encephalomyocarditis (EMCV) (31).

### IFN-I has reduced anti-WNV activity in TRIM6-deficient cells

Next, we sought to evaluate further the impact of TRIM6 on the antiviral efficiency of IFN-I against WNV. Prior to infection, wt and TRIM6-KO A549 cells were treated with 100U of recombinant human IFNβ for 4 hours prior to infection with WNV (MOI 5.0) for 24 hours (Figure 2). Pre-treatment with IFNβ decreased viral load in both wt and TRIM6-KO cells, however IFN-I pre-treatment was significantly less effective in inhibiting WNV replication in TRIM6-KO (40 fold) as compared to wt controls (63 fold). As expected, this result indicates that IFN-I signaling is suboptimal in the absence of TRIM6, enabling WNV to replicate to higher titers, and suggests that expression of TRIM6-dependent ISGs may be involved in establishing an optimal IFN-I mediated anti-WNV response.

**Figure 2.**
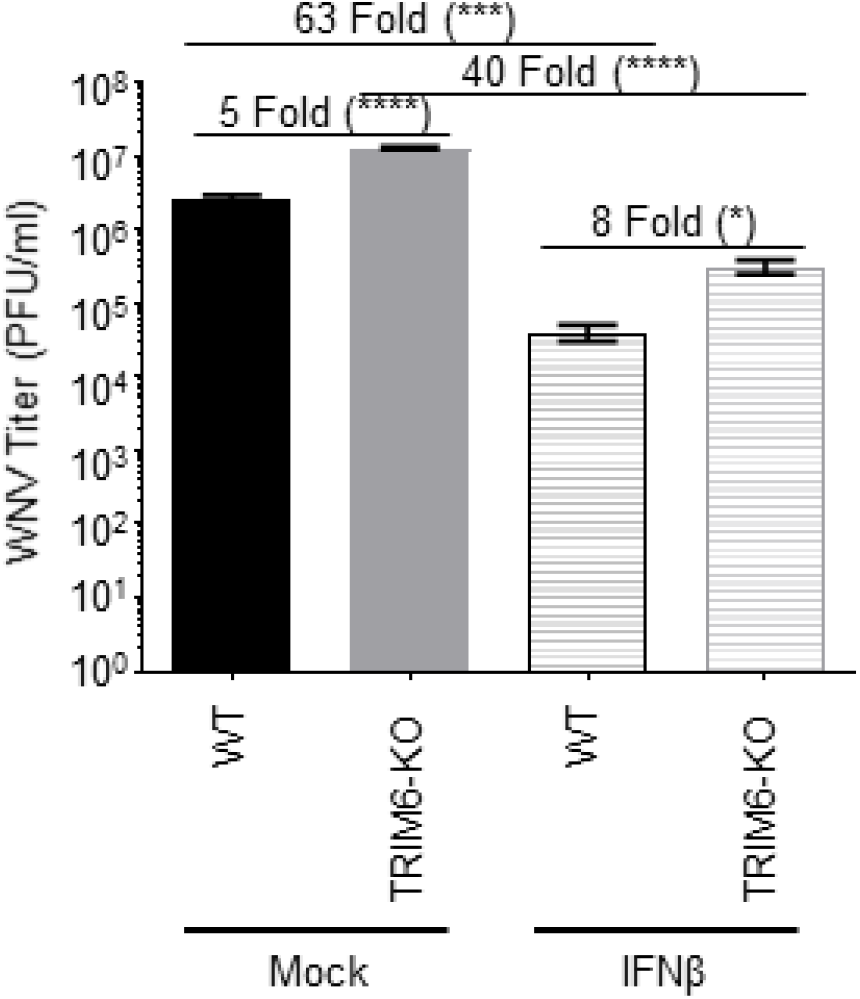
IFN-I Pre-treatment is Less Efficient in Antagonizing WNV Replication in TRIM6-KO Cells. wt or TRIM6-KO cells were treated with recombinant human IFNβ-1a (100U) for 4 hours prior to infection with WNV 385-99 (MOI 5.0) for 24 hours. Supernatants from infected cells were titrated and viral load was calculated via plaque assay. Error bars represent standard deviation (n=3). One-way ANOVA with Tukey’s post-test was performed to assess statistical significance; ****p <0.0001, *p <0.05. Fold change reported in parenthesis. All experiments were performed in triplicate.

### VAMP8 is induced in a TRIM6-dependent manner

To identify other genes affected as a consequence of TRIM6’s absence, next generation sequencing (NGS) of mock (Figure 3A) and WNV-infected (Figure 3B) wt and TRIM6-KO cells was performed. During WNV infection, canonical ISGs were identified as being expressed at lower levels in the TRIM6-KO compared to wt cells, which validates the methodology (Figure 3B). Several canonical ISGs down-regulated in TRIM6-KO cells have previously been described to inhibit WNV replication, including *Ifitm2* (42) and *Ifitm3* (12), or their loss of function is associated with increased WNV susceptibility, including *Mx1* (21, 43) and *OasL* (43). We elected to investigate other genes not previously described to regulate WNV replication or IFN-I pathways. One of the strongest downregulated genes in both mock and infected cells, VAMP8, was chosen as a target for further mechanistic validation (Figure 3A-B). Although VAMP8 has previously been noted to play an antiviral role in response to Japanese encephalitis virus (JEV) (44), the mechanism has not been reported. Further, VAMP8’s described roles are diverse, but a connection to the antiviral IFN-I pathway has not been described.

**Figure 3.**
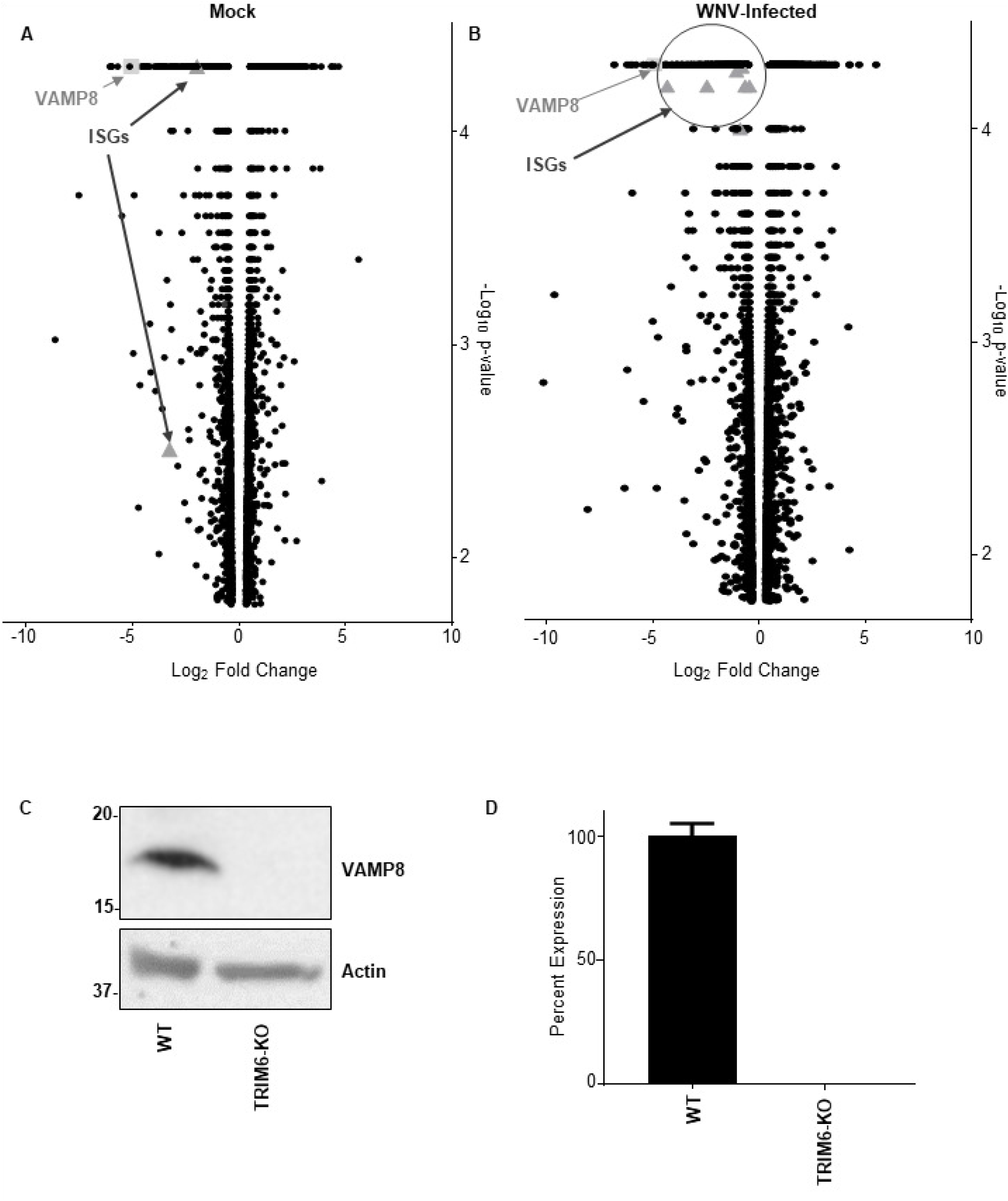
Transcription of Canonical Interferon-Stimulated Genes and VAMP8 are Down-Regulated in TRIM6 Knockout Cells. Transcriptional profiling of cellular mRNA by Next Generation Sequencing of mock (A) or WNV 385-99 infected (MOI 5.0) (B) wt or TRIM6-KO A549 at 24 hours post-infection. Log_2_ fold change was calculated as TRIM6-KO/mock with genes down-regulated in TRIM6-KO cells on the left (negative values) and up-regulated in TRIM6-KO cells on the right (positive values). The –log_10_ p-value represents the significance. VAMP8 data point is represented as a light grey square and interferon-simulated genes (ISGs) are represented as dark grey triangles. Validation of VAMP8 expression at the protein (immunoblot in C) or RNA (RT-qPCR in D) levels in wt or TRIM6-KO cells. Error bars represent standard deviation (n=3) and VAMP8 expression validation experiments were performed in triplicate and repeated three times.

After confirming lower expression of VAMP8 at the translational (Figure 3C) and transcriptional (Figure 3D) levels in TRIM6-KO cells, the role of VAMP8 in regulating WNV replication was interrogated. Therefore, wt A549 cells were transfected with a VAMP8-targeting siRNA pool or non-targeting control siRNA (ntc) for 24 hours prior to WNV infection (MOI 0.1). VAMP8 knockdown (kd) had no measurable effect on WNV replication (Figure 4A), suggesting that VAMP8 does not have a direct anti-WNV function. VAMP8 knockdown was validated by western blot, showing undetectable levels of protein, with a clear upregulation in VAMP8 protein by 24 hpi in ntc transfected cells (Figure 4B). Upon infection, phosphorylation of IRF3 in VAMP8-kd cells was not significantly different as compared to ntc cells, suggesting that VAMP8 is not involved in the IFN-I induction arm of the pathway (Figures 4B, S2A). In contrast, the amount of pSTAT1 (Y701) was notably lower in the VAMP8-kd cells, suggesting impairment in the IFN-I signaling pathway (Figures 4B, S2B).

**Figure 4.**
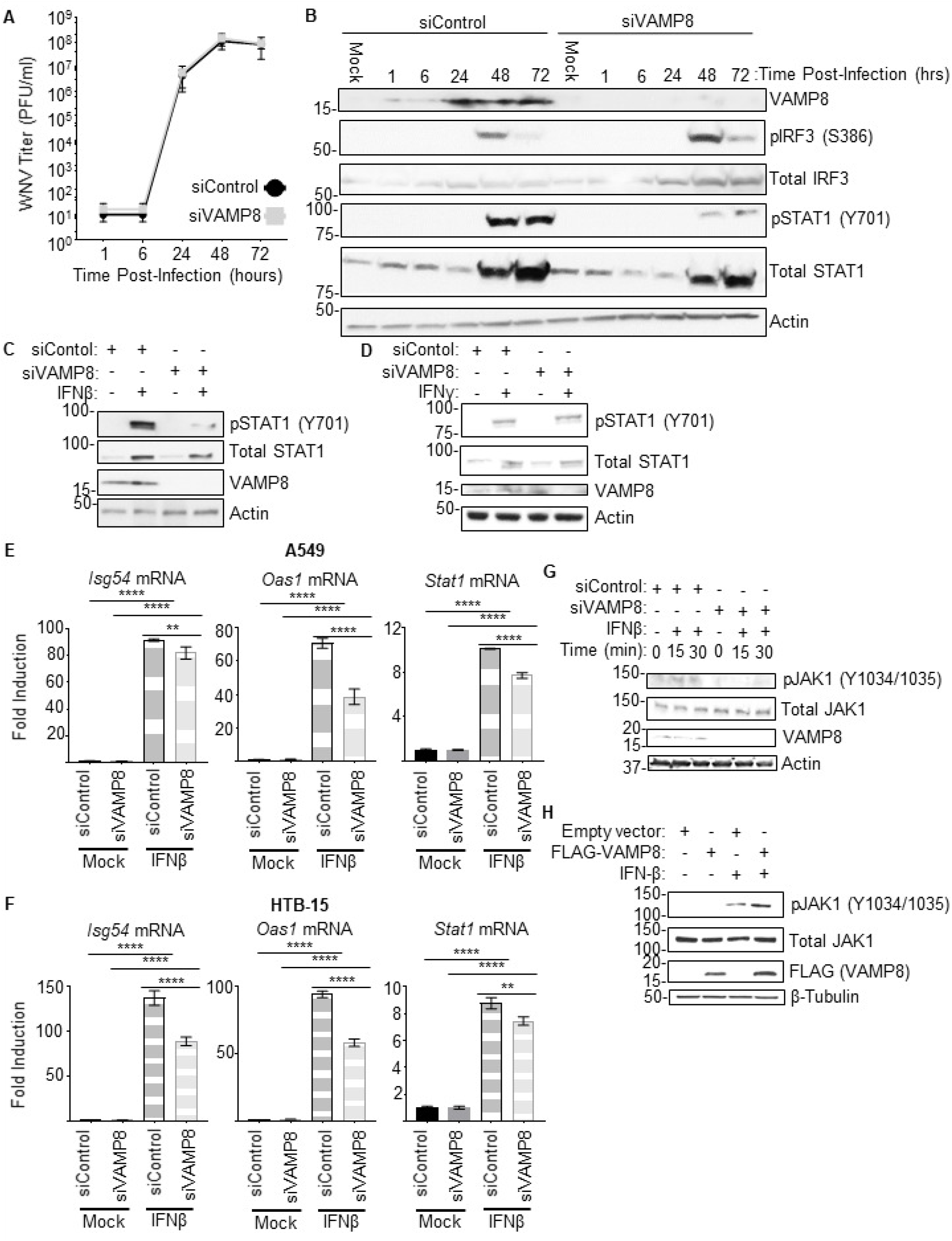
Depletion of VAMP8 Impairs IFN-I Signaling but does not alter WNV Replication. Wild-type (wt) A549 cells were treated with non-targeting control (control) or VAMP8-targeting (VAMP8) siRNAs for 24 hours followed by infection with WNV 385-99 (MOI 0.1) for 72 hours (A, B). Supernatants and lysates of infected cells were collected at 1, 6, 24, 48, and 72 hours post-infection to assess viral load by plaque assay (A) and protein expression and phosphorylation by western blot (B). wt A549 (C, D, E, G) or HTB-15 (F) cells were treated with non-targeting control (siControl) or VAMP8-targeting (siVAMP8) siRNAs for 24 hours followed by treatment with recombinant human IFNβ-1a (500U/mL) (C, E, F, G) or human IFNγ (500U/mL) (D). IFN treatments shown in C, D, E, F, were for 16 hr. Cells were lysed and either protein (B,C, D, G, H) or RNA (E,F) was isolated for analysis by western blot or qRT-PCR, respectively. FLAG-tagged VAMP8 or empty vector was transfected into HEK293T cells for 24 hours prior to treatment with human IFNβ-1a (1000U/mL) for 1 hour and protein lysates were collected to assess Jak1 activation (H). Error bars represent standard deviation. Gene expression data was analyzed using one-way ANOVA with Tukey’s post-test to assess statistical significance (E,F); p**** <0.0001, p** <0.01. No statistical significance found in (A). Experiment performed in triplicate.

To further confirm whether VAMP8 is involved in regulation of the IFN-I signaling pathway in the absence of virus infection, wt A549 cells were transfected with VAMP8-targeting or ntc siRNAs followed by treatment with IFNβ for 16 hours. As expected, total STAT1 was induced in both VAMP8-and ntc-siRNA treated cells following IFNβ stimulation, but the level of total STAT1 in VAMP8-kd cells was slightly attenuated (Figure 4C). VAMP8’s effect on STAT1 activation is more evident; however, with a strong reduction in the amount of pSTAT1 (Y701) (Figures 4C, S2C). To evaluate whether the observed impairment in pSTAT1 (Y701) is specific to type-I IFN signaling, we treated ntc- and VAMP8-kd cells with the type II interferon (IFNγ) for 16 hours. In contrast to the observed defect in response to IFNβ, there was no detectable difference in pSTAT (Y701) between siControl and siVAMP8-treated cells following IFNγ stimulation (Figures 4D, S2D). Consistent with a potential role of VAMP8 in regulating STAT1 phosphorylation downstream of IFN-I signaling, mRNA expression levels upon IFNβ treatment of ISGs including *Isg54*, *Oas1*, and *Stat1* were significantly reduced in VAMP8-kd as compared to controls in both A549 (Figure 4E) and a brain-derived cell line (glioblastoma HTB-15 cells) (Figure 4F).

In an effort to determine whether VAMP8 functions at the level of STAT1 or upstream in the pathway, we assessed the effects of VAMP8 on Jak1 activation, which is responsible for phosphorylation of STAT1 on Y701. In the absence of VAMP8, phosphorylation of Jak1 (pY1034/1035) was reduced compared to control cells (Figures 4G, S2E). Further, ectopic expression of FLAG-tagged VAMP8 in HEK293T cells enhanced pJak1 following IFNβ stimulation compared to empty vector-transfected cells (Figure 4H, S2F). Overall, the above evidence supports that 1) VAMP8 expression is TRIM6-dependent, 2) VAMP8 does not directly affect WNV replication, and 3) VAMP8 is involved in positive regulation of IFN-I signaling on or upstream of Jak1 phosphorylation.

### VAMP8 Knockdown Enhances WNV Replication in Cells Pre-treated with Type I IFN

Since VAMP8 modulates the IFN-I system, but does not appear to alter WNV replication, we examined whether exogenous IFN-I pre-treatment would reveal a functional defect in IFN-I signaling in VAMP8-kd cells during WNV infection. Prior to infection, wt A549 or brain-derived HTB-15 cells were treated with siRNA (VAMP8 or ntc) followed by IFNβ treatment. Although IFNβ pre-treatment reduced viral production in both groups, it was less efficient in protecting VAMP8-kd cells against WNV replication as compared to ntc-cells (A549: ntc siRNA: 508 fold; VAMP8 siRNA: 79 fold (Figure 5A); HTB-15: ntc siRNA: 430 fold; VAMP8 siRNA: 20 fold (Figure 5B)). As opposed to previous experiments showing no impact on WNV replication following VAMP8-kd, the combination of IFN-β pre-treatment and VAMP8 siRNA showed an 8-fold (A549) (Figure 5A) or 22-fold (HTB-15) (Figure 5B) increase in the replication of WNV over cells treated with ntc siRNA and IFNβ. This result suggests that VAMP8 plays a functional role in IFN-I signaling during WNV infection.

**Figure 5.**
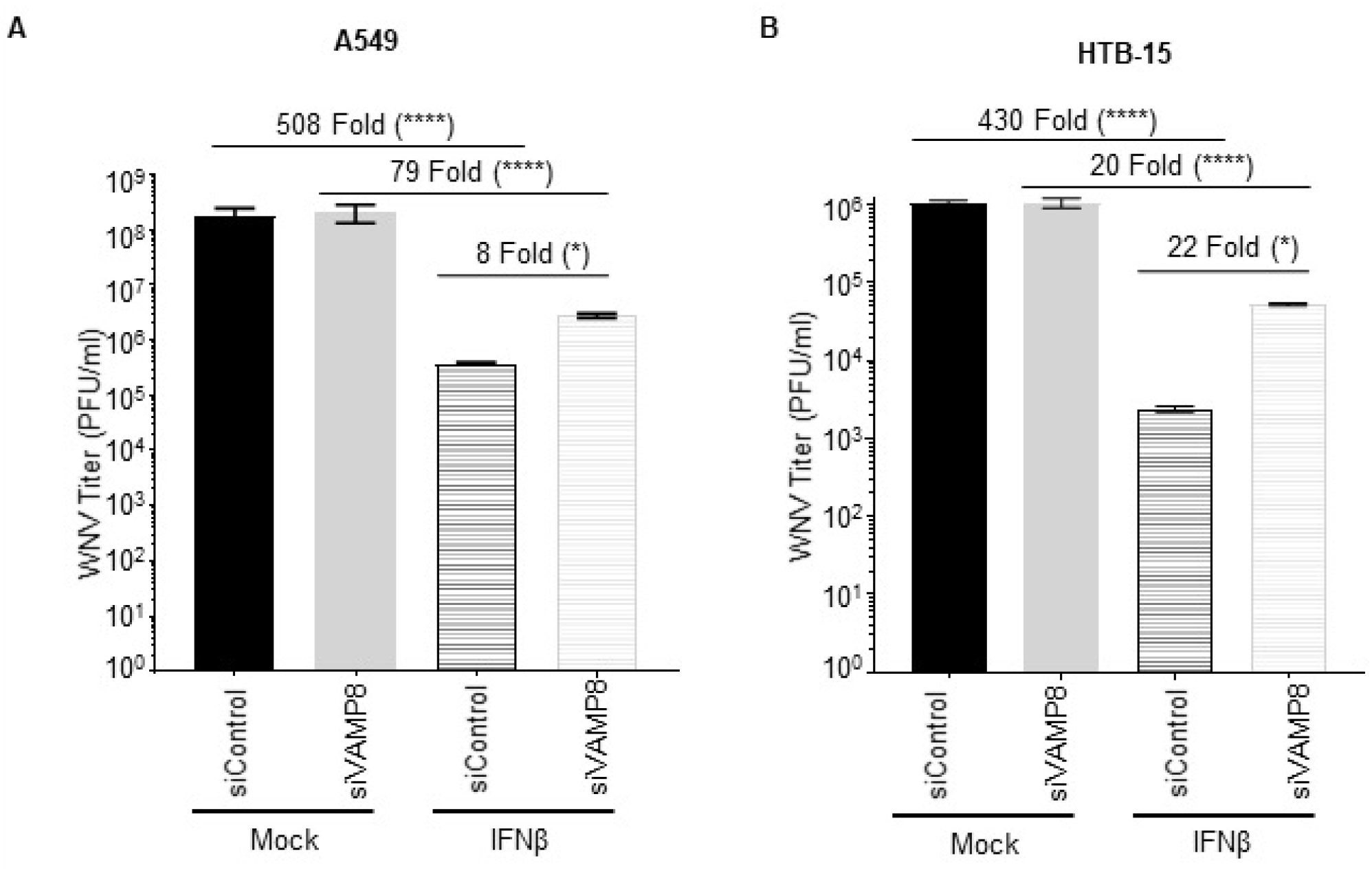
VAMP8 is Important for efficient establishment of an anti-WNV response mediated by IFNβ. wt A549 (**A**) or HTB-15 (**B**) cells were treated with non-targeting control (siControl) or VAMP8-targeting (siVAMP8) siRNAs for 24 hours then treated with recombinant human IFNβ-1a (500U/mL) for 16 hours prior to infection with WNV 385-99 (MOI 5.0) for 24 hours. Supernatants from infected cells were titrated and viral load was calculated via plaque assay. Error bars represent standard deviation. One-way ANOVA with Tukey’s post-test was performed to assess statistical significance; ****p <0.0001, *p <0.05. Fold change reported in parenthesis. Experiment completed in triplicate.

### VAMP8 Over-expression Attenuates WNV Replication

To further investigate whether VAMP8 overexpression can induce an anti-WNV response and whether this requires the presence of TRIM6, we transfected wt or TRIM6-KO A549 cells with either empty vector (pCAGGS) or FLAG-tagged VAMP8 (FLAG-VAMP8) 30 hours prior to infection with WNV (MOI 5.0). As predicted, in wt A549 cells, the viral load was significantly decreased (8.5 fold) in cells over-expressing FLAG-VAMP8 compared to empty vector control (Figure 6A). In contrast, when FLAG-VAMP8 was over-expressed in TRIM6-KO cells the viral load was not significantly different from cells transfected with empty vector (Figure 6A). This result suggests that VAMP8’s attenuation of WNV replication requires TRIM6 activity and can be explained by VAMP8’s inability to compensate for the defect in pSTAT1 (S708) in TRIM6-KO cells (Figure 6B, S3A). This phosphorylation on S708 of STAT1 is required for stabilization of the ISGF3 complex (41) and induction of the complete set of ISGs (30), including ISGs known to be involved in the anti-WNV response (11, 13, 42). Therefore, while VAMP8 is responsible for promoting STAT1-Y701 phosphorylation, which can only account for limited antiviral effects, induction of the complete set of anti-WNV ISGs does not occur unless the TRIM6-IKKε-STAT1-S708 arm of the pathways is active. To rule out a potential role of TRIM6 in VAMP8-mediated promotion of STAT1-Y701 phosphorylation, we over-expressed FLAG-VAMP8 in wt or TRIM6-KO A549 cells prior to treatment with IFNβ for 16 hours. Unexpectedly, over-expression of VAMP8 induced pSTAT1 (Y701) following IFNβ stimulation in wt but not TRIM6-KO A549 cells (Figure 6C, S3B), suggesting that TRIM6 is not only required for VAMP8 expression (as described above in Figure 3), but also for VAMP8 activity. To further test this possibility, we investigated whether TRIM6 interacts with VAMP8. Co-immunoprecipitation assays (coIP) showed that FLAG-VAMP8 co-precipitated HA-TRIM6 (Figure 6D) and HA-TRIM6 co-precipitated FLAG-VAMP8 in the reverse coIP (Figure 6E). Taken together these data suggest that TRIM6 is important for VAMP8 expression levels as well as VAMP8 activity downstream the IFN-I receptor and for an optimal anti-WNV response.

**Figure 6.**
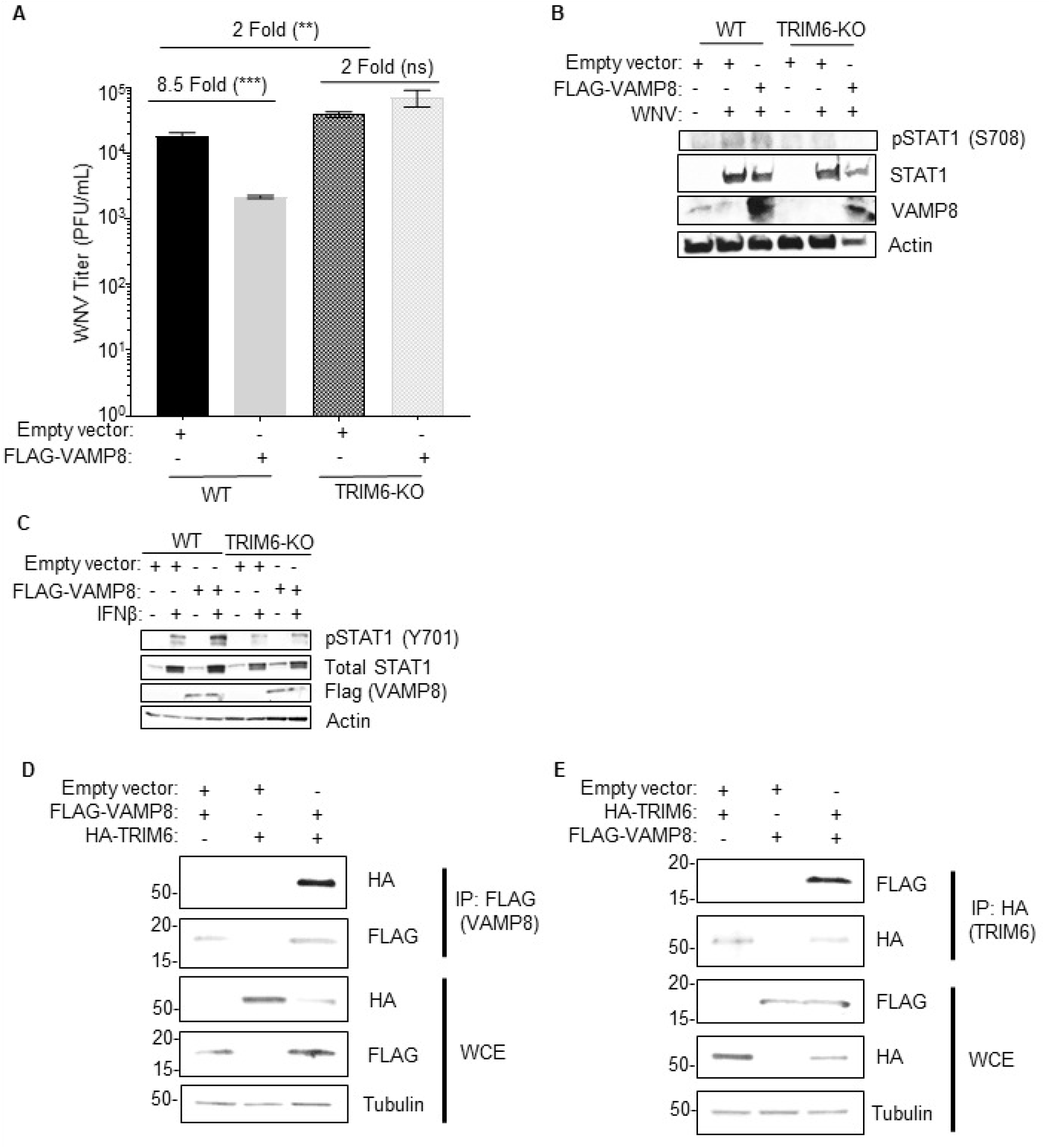
VAMP8 Over-expression Attenuates WNV Replication. Wild-type or TRIM6-KO A549 cells were transfected with 250ng of empty vector or Flag-VAMP8 for 30 hours then infected with WNV 385-99 at a MOI 5.0 for 24 hours (A) or treated with IFNβ (500U/mL) for 16 hours. Supernatants from infected cells were titrated and viral load was calculated via plaque assay (A). Protein lysates from were collected to measure STAT1 activation (p S708) (B) or (p Y701) (C) and to confirm equal levels of VAMP8 overexpression in wt and TRIM6-KO cells. FLAG-tagged VAMP8 was co-transfected with empty vector or HA-tagged TRIM6 into HEK293T cells for 24 hours (D, E). Protein lysates from the co-transfected cells were immunoprecipitated (IP) with either anti-FLAG (D) or anti-HA (E) beads overnight prior. Whole-cell lysates (WCE) and IP samples were immunoblotted to assess expression and VAMP8-TRIM6 interaction. Error bars represent standard deviation. A student’s t-test was used to assess statistical significance; ***p <0.001, **p<0.01, ns no significance. Experiment performed in triplicate.

## DISCUSSION

Our study demonstrates the relevance of TRIM6 in regulating the IFN-I pathway during WNV infection and identifies VAMP8 as a factor functionally involved in IFN-I signaling. Extensive research has implicated the TRIM family of proteins in both regulation of the innate immune response and viral replication (45–50). Specifically, TRIM6 has been shown to facilitate the formation of unanchored K48-linked polyubiquitin chains that provide a scaffold for IKKε binding, ultimately resulting in IKKε activation and STAT1 phosphorylation at S708 (31). Phosphorylation of STAT1 at S708 is important to sustain IFN-I signaling and to express a unique subset of ISGs (30, 41). The relevance of IKKε-dependent gene expression has previously been described for WNV, and in the absence of ISG54 or IKKε mice have an increased susceptibility (11). These experiments served as a rationale for exploring the functional role of TRIM6 during WNV infection.

As expected, we observed an increase in WNV replication in TRIM6-KO cells in parallel with attenuated TRIM6-dependent activation of IKKε (T501 phosphorylation), IKKε-dependent STAT1 S708 phosphorylation, and IKKε-dependent gene expression. There was impaired *Ifnb* and *Isg54* mRNA induction in TRIM6-KO cells at 6 and 24 hpi, respectively, but higher levels of induction at 72 hpi. The transient effect of TRIM6 on the IFN-I pathway may be related to the increase in pTBK1 (S172) in TRIM6-KO cells compensating for the impaired IKKε-dependent response to WNV infection. Another possible explanation is that the ISGs higher in TRIM6-KO cells are either pSTAT1 (S708) independent (STAT1 and IRF7, (30)) or can be induced directly via IRF3 or IRF7 by binding their promoters (e.g. ISG54,(51)), while Oas1 is a TRIM6-IKKε-pSTAT1 (S708)-dependent ISG (30, 31), which is absent in TRIM6-KO cells. Further, redundant or alternative pathways not investigated here may be enacted in the TRIM6-KO cells to control WNV replication. Although no other factor has been shown to synthesize the unanchored K48-polyubiquitin chains required for IKKε activation, we cannot exclude that other TRIM members or other E3-Ub ligases may compensate for the loss of TRIM6. Alternatively, TRIM6 may play important roles in other pathways (i.e. NF-ĸB), resulting in cytokine dysregulation and/or induction of IFNβ by TRIM6-independent pathways. Further, emergent WNV strains encode a functional 2-O methylase in their non-structural protein 5 that prohibits IFIT proteins, specifically murine ISG54 and human ISG58, from suppressing viral mRNA expression (52). Since WNV antagonizes components of this pathway, we cannot rule out the possibility that WNV proteins target TRIM6 to impede IKKε-dependent expression of WNV-restricting ISGs. For example, the matrix protein of Nipah virus (family *Paramyxoviridae*) works to promote the degradation of TRIM6 during viral infection to promote viral replication through impaired IKKε signaling and thus a blunted IFN-I response (46). WNV protein antagonism of TRIM6 could also preclude observing more severe differences in WNV replication between wt and TRIM6-KO cells. Alternatively, TRIM6 could play an essential role in IKKε-dependent signaling but could also be hijacked by a virus to facilitate viral replication. In a previous study, we showed that TRIM6 directly promotes the replication of ebolavirus (family *Filoviridae*) through interactions with VP35 and that VP35 antagonizes TRIM6’s capacity to promote IFN-I signaling (45).

Importantly, although the experiments presented here were performed with cell lines, our previous published studies suggest that TRIM6 plays an important antiviral role via the IFN-I system in primary human monocyte derived dendritic cells (hMDDC) and the antiviral response to two other viruses, including SeV and EMCV, and to influenza virus in lungs of mice (31). Therefore, the effects of TRIM6 are not restricted to cell line or to WNV.

Although we identified several ISGs differentially expressed in TRIM6-KO compared to wt cells following WNV infection, we also identified *Vamp8* to be significantly down-regulated under both basal conditions and WNV infection. VAMP8 has not been previously described to affect WNV replication or the IFN-I pathway. Although its role was not described, VAMP8 had been identified as an antiviral factor in an siRNA screen for JEV, another mosquito-borne flavivirus (44). Here we showed, in contrast to that seen with JEV, VAMP8 knockdown does not directly affect WNV replication. The impairment of STAT1 Y701 phosphorylation during WNV in the VAMP8-kd cells lent evidence that VAMP8 could be involved in IFN-I signaling. Since WNV efficiently impairs IFN-I induction until nearly 24 hpi, VAMP8 depletion may not substantially impede of IFN-I signaling during WNV infection in a tissue culture system. Evaluation of VAMP8’s role in IFN-I signaling in the absence of WNV infection revealed a striking impairment in STAT1 Y701 phosphorylation and a modest inhibition of ISG gene expression in VAMP8-depleted cells. The impairment of STAT1 activation (pY701) in the absence of VAMP8 occurs specifically downstream of IFN-I but not IFN-II signaling, and a delay and reduction of Jak1 activation may contribute to this phenotype. Further, following IFNβ treatment, VAMP8-knockdown less efficiently inhibited WNV replication, which provides support that VAMP8 mediates a functional step in the IFN-I signaling pathway. We also showed that VAMP8 over-expression modestly inhibits WNV replication in wt but not TRIM6-KO cells, which suggest that both VAMP8 and TRIM6 are necessary for induction of the full set of antiviral ISGs. There are two possible explanations for these results; a) VAMP8 is important for efficient phosphorylation on STAT1-Y701 (independent of STAT1-S708 phosphorylation by the TRIM6-IKKε arm), and in parallel the TRIM6-IKKε arm is required for phosphorylation on STAT1-S708. Both of these events are necessary for efficient induction of the full set of ISGs and antiviral activity, or b) TRIM6 could be important for directly regulating VAMP8-mediated JAK1-STAT1-Y701 phosphorylation and activation, in addition to promoting IKKε-dependent STAT1-S708 phosphorylation. We favor the latter explanation because VAMP8 requires the presence of TRIM6 to attenuate WNV replication (Figure 6A) and to induce IFN-dependent STAT1-Y701 phosphorylation (Figure 6C). In addition, TRIM6 interacts with VAMP8 (Fig 6D,E), and TRIM6 deficiency impedes the induction of STAT1-S708 phosphorylation independent of VAMP8 expression (Fig 6B).

Although our results suggest that TRIM6 and VAMP8 interact and VAMP8’s regulation of the IFN-I pathway is TRIM6-depedent, the detailed mechanism of how VAMP8 promotes JAK1 activation downstream the IFN-I receptor is unknown. VAMP8 is involved in endocytosis (32), vesicle-vesicle fusion (35), and exocytosis (33, 34, 36) in various cell types including leukocytes (37, 40), and various secretory cells (33, 34, 36) including human lung goblet cells. As a vesicular SNARE (v-SNARE), VAMP8 on the surface of a vesicle interacts with SNAREs on the target membrane surface to facilitate membrane fusion (33, 35, 37). Potential mechanisms of the IFN-I pathway, which VAMP8 may regulate, include surface expression of the IFNAR receptor or recycling of receptor components to the plasma membrane. Although we did not find interaction of VAMP8 with the IFNAR or JAK1 (data not shown), it is still possible that VAMP8 could regulate JAK1 activity or IFNAR function. VAMP8 has been described to regulate the surface expression of a water transport channel, aquaporin 2, in the kidney (39). Despite the reduced surface expression, the total amount of aquaporin 2 is higher in the cell, but it is retained in vesicles below the plasma membrane (39). Alternatively, VAMP8 might influence the secretion of factors or the oxidative condition of the microenvironment important to maintain IFN-I signaling. In phagocytic cells infected with *Leishmania*, VAMP8 regulates the transport of NADPH oxidase to the phagosome to facilitate optimal conditions for peptide loading into MHC class I molecules (40). Although VAMP8 would not be regulating phagocytosis in this model of WNV infection, it is possible that VAMP8 regulates NADPH oxidase localization affecting the oxidative environment of the infected cell and consequently IFN-I signaling (53–55). TRIM6 may either directly affect VAMP8 expression or may act indirectly through a yet unidentified secondary factor. Knowing that TRIM6 is important in regulating IKKε activation and function (31), we assessed whether IKKε alters VAMP8 expression. There was no difference in VAMP8 expression between IKKε-wt and -KO murine embryonic fibroblasts under basal conditions (Figure S4A). We did identify two transcription factors, HNF4α and HNF4γ, with binding sites upstream of VAMP8’s transcription initiation site and within a VAMP8 intron that have significantly lower gene expression in TRIM6-KO compared to wt A549 cells (Figure S4B-D). The expression of wt TRIM6, but not the TRIM6 catalytic mutant (C15A), partially rescues the expression of VAMP8 in TRIM6-KO A549 cells suggesting the presence of TRIM6 and its ubiquitin ligase activity are required for VAMP8 expression (Figure S4E). Our study indicates a new role for VAMP8 in the IFN-I pathway, which is regulated by TRIM6 during viral infection (Figure 7).

**Figure 7.**
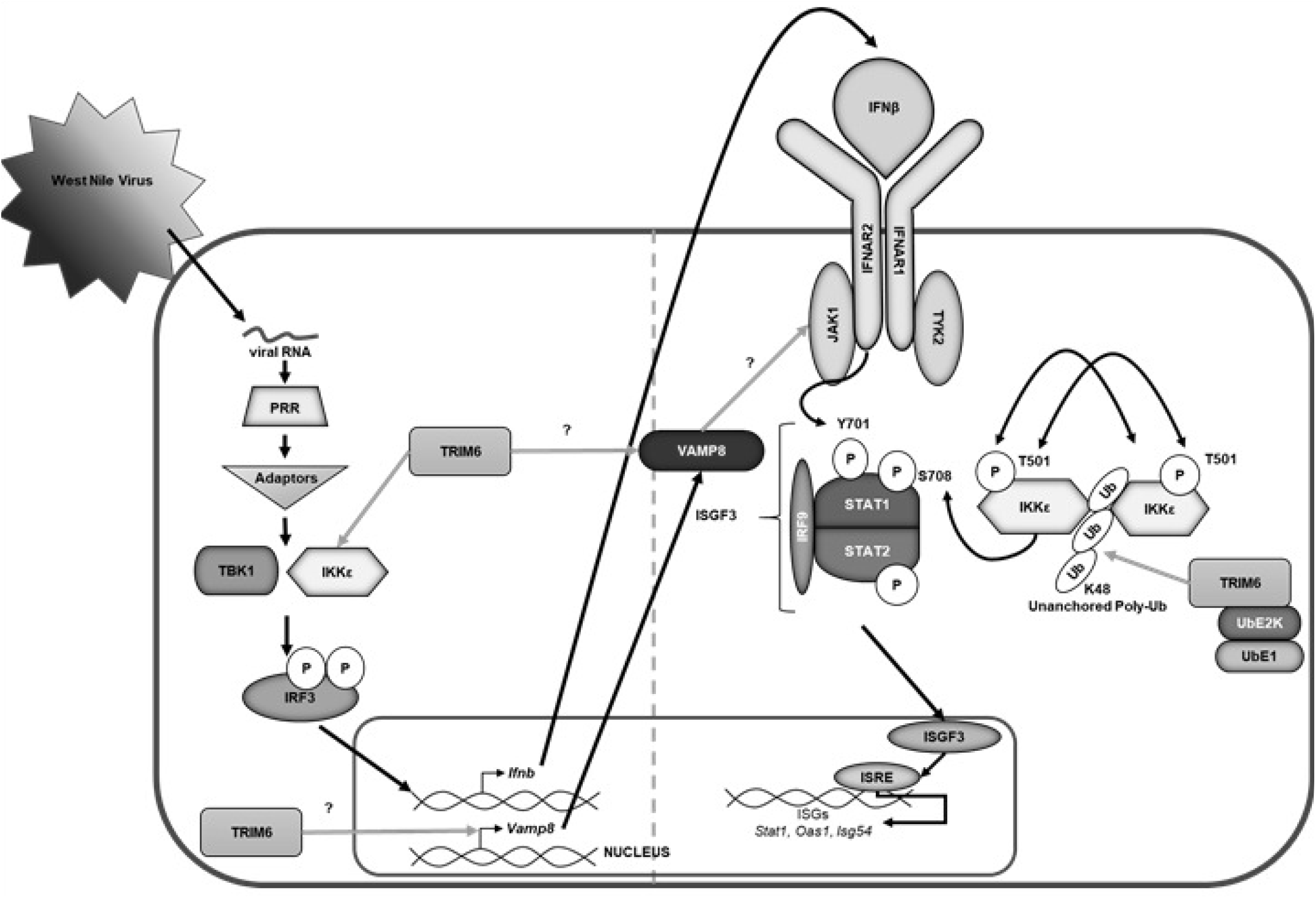
Graphical Summary. Following virus infection, viral RNA is recognized by pathogen recognition receptors (PRRs). PRRs then signal through their adaptors, triggering the activation of kinases TBK1 and IKKε, which phosphorylate and activate the transcription factor IRF3. Once activated, IRF3 translocates to the nucleus and, in concert with other factors not indicated, promotes the transcription of IFNβ. IFNβ is then secreted and signals in an autocrine or paracrine manner through the type I IFN receptor (IFNAR). The kinases (Jak1 and Tyk2) associated with IFNAR then facilitate the phosphorylation of STAT1 at tyrosine (Y) 701 and STAT2 in an IKKε-independent manner. Phosphorylated STAT1 and STAT2 interact with IRF9 to form the ISGF3 complex, which translocates to the nucleus to promote the transcription of genes with interferon stimulated response elements (ISRE) including *Stat1, Oas1,* and *Isg54*. In addition to IKKε independent IFN-I signaling, the E3 ubiquitin ligase TRIM6 facilitates IKKε-dependent IFN-I signaling. TRIM6, in coordination with the ubiquitin activating (UbE1) and ligating (UbE2K) enzymes to facilitate the formation of K48-linked unanchored poly-ubiquitin chains, which act as a scaffold for the oligomerization and cross-phosphorylation of IKKε at threonine (T) 501 (30). TRIM6 also facilitates activation of IKKε during IFN-I induction. During IFN-I signaling, activated IKKε phosphorylates STAT1 at serine (S) 708. STAT1 phosphorylation at S708, an IKKε-dependent modification, facilitates the formation of an ISGF3 complex with different biophysiological properties which allows the ISGF3 complex to have enhanced binding to certain ISRE-containing promoters ultimately inducing the complete ISG profile. When STAT1 is phosphorylated only at Y701 (in the absence of IKKε and/or TRIM6), IFN-I signaling results in induction of different and incomplete ISG profile (30, 31, 41). Although the mechanism is currently unknown (question mark), TRIM6 induces VAMP8 expression and VAMP8 activity. VAMP8, in turn, is important for the optimal activation of Jak1 and subsequently STAT1 (Y701) required for an efficient antiviral response.

Elucidating the interactions of the human immune system with viral infection is essential to understanding viral pathology, as well as identifying cellular targets for antiviral drug development. Our work has identified a novel IFN-I-related host factor that is important in the regulation of WNV replication and in the life cycles of other viruses. This may provide a conserved target for the development of anti-viral strategies and for the elucidation of further conserved pathways in host-pathogen interaction.

## MATERIALS & METHODS

### Viruses & Cells

West Nile Virus (WNV) isolate 385-99 was obtained from the World Reference Center for Emerging Viruses & Arboviruses (UTMB, Dr. Robert Tesh). A549, HEK 293T, HTB-15 (U-118MG), and CCL-81 lines were obtained from the American Type Culture Collection. Wild-type and IKBKE^-/-^ murine embryonic fibroblasts were a kind gift from Dr. tenOever at Ichan School of Medicine at Mount Sinai, and were described previously (30, 31). TRIM6 knockout cells were prepared as previously described (45). All lines were maintained in DMEM (Gibco), supplemented with 10% Fetal Bovine Serum (Atlanta Biologicals). Infections were performed in DMEM supplemented with 2% Fetal Bovine Serum (FBS), and 1% Penicillin/Streptomycin (Gibco). For growth kinetics experiments, 150,000 wt or TRIM6-KO A549 or HTB-15 cells/well were infected with 100μL of WNV multiplicity of infection (MOI) 0.1 or 5.0 for 1 hour at 37°C, 5% CO_2_ then the inoculum was removed and washed 3 times with 1mL of 1X PBS. After the cells were washed, 1mL of DMEM supplemented with 2% FBS was added to each well. Supernatant (150μL) was collected at the designated time points for plaque assay. Plaque assays were performed in 12-well plates containing 200,000 CCL-81 cells/well. Viral samples were diluted log fold and applied to the monolayer. Following 1 hour in a humidified 37°C, 5% CO_2_ incubator, semisolid overlay containing MEM, 2% Fetal Bovine Serum, 1% Penicillin/Streptomycin and 0.8% Tragacanth (Sigma Aldrich) was applied. Overlay was removed after 72 hours, and monolayers were fixed and stained with 10% Neutral Buffered Formalin (Thermo Fisher Scientific) and Crystal Violet (Sigma Aldrich). Plaques were enumerated by counting and graphed. All manipulations of infectious West Nile Virus were performed in Biological Safely Level 3 facilities at UTMB.

### IFN Treatment

Cells were treated with either 100U (wt *vs* TRIM6-KO) or 500U (VAMP8) of recombinant human IFNβ-1a (PBL Assay Science) or 500U/mL of IFNγ (PBL Interferon Source) for either 15 and 30 minutes to assess Jak1 activation, 16 hours to investigate STAT1 activation, and 4 hours or 16 hours prior to WNV infection.

### RNA Isolation and qRT-PCR

At the indicated timepoint per experiment, media was removed from the cell monolayer, and 1mL Trizol Reagent (Thermo Fisher Scientific) was added. RNA was isolated using Zymo Direct-zol RNA Miniprep Kits as per manufacturer instruction with in-column DNase treatment. Isolated RNA was then reverse transcribed using the high capacity cDNA reverse transcription kit (Applied Biosystems). The cDNA was then diluted 1:3 in nuclease-free water (Corning). Relative gene expression (primers listed in Supplementary Table 1) was determined using the iTaq^TM^ Universal SYBR green (Bio-Rad) with the CFX384 instrument (Bio-Rad). The relative mRNA expression levels were analyzed using the CFX Manager software (Bio-Rad). The change in threshold cycle (ΔCT) was calculated with 18S gene served as the reference mRNA for normalization. When indicated, fold change was calculated by dividing the −2^Δ*Ct*^ value for the treated sample by its respective mock.

### RNA Sequencing & Analysis

A549 (wt and TRIM6KO) cells were infected at a high MOI (5.0) and RNA isolated 24 hours post infection. RNA quality was assessed by visualization of 18S and 28S RNA bands using an Agilent BioAnalyzer 2100 (Agilent Technologies, CA); the electropherograms were used to calculate the 28S/18S ratio and the RNA Integrity Number. Poly-A+ RNA was enriched from total RNA (1 μg) using oligo dT-attached magnetic beads. First and second strand synthesis, adapter ligation and amplification of the library were performed using the Illumina TruSeq RNA Sample Preparation kit as recommended by the manufacturer (Illumina, Inc). Library quality was evaluated using an Agilent DNA-1000 chip on an Agilent 2100 Bioanalyzer. Quantification of library DNA templates was performed using qPCR and a known-size reference standard. Cluster formation of the library DNA templates was performed using the TruSeq PE Cluster Kit v3 (Illumina) and the Illumina cBot workstation using conditions recommended by the manufacturer. Paired end 50 base sequencing by synthesis was performed using TruSeq SBS kit v3 (Illumina) on an Illumina HiSeq 1500 using protocols defined by the manufacturer. The alignment of NGS sequence reads was performed using the Spliced Transcript Alignment to a Reference (STAR) program, version 2.5.1b, using default parameters (56). We used the human hg38 assembly as a reference with the UCSC gene annotation file; both downloaded from the Illumina iGenomes website. The –quantMode GeneCounts option of STAR provided read counts per gene, which were input into the DESeq2 (version 1.12.1) (57) differential expression analysis program to determine expression levels and differentially expressed genes.

### Transfections and Immunoprecipitations

Transient knockdown of endogenous VAMP8 in wt A549 was done in 12-well plates. Briefly, 20 pmol of Smartpool ON-TARGETplus Non-targeting (D-001810-10-05) or ON-TARGETplus VAMP8 (L-013503-00-0005) siRNA (Dharmacon) were transfected with Lipofectamine RNAiMAX (Invitrogen) following the manufacturer’s instructions. Cells were transfected with siRNA 24 hours prior to infection or IFNβ treatment. The efficiency of VAMP8 knockdown was monitored by qRT-PCR or western blot. Wild-type or TRIM6-KO A549 or HEK 293T cells were transfected with 250ng of pCAGGS (empty vector) or FLAG-tagged VAMP8 (Origene) using Lipofectamine 3000 (Invitrogen) according to the manufacturer’s instructions. Cells were transfected with plasmid DNA 30 hours prior to WNV infection or 24 hours prior to human IFNβ-1a treatment. For immunoprecipitations, cells were lysed in RIPA complete buffer and centrifuged at 15,000 rpm at 4°C for 20 minutes to clarify the lysates. The clarified lysates were incubated with anti-HA or anti-FLAG coated beads (Sigma) overnight at 4°C. The beads were washed with RIPA (NEM + IA) seven times then resuspended in 2X laemmli with 5.0% beta-mercaptoethanol and boiled at 100°C for 10 minutes.

### Western Blotting

Infected or IFNβ-treated cells were lysed in 2X Laemmli buffer with β-ME and boiled at 100°C for 10 minutes prior to removal from BSL-3. For immunoblotting, proteins were resolved using SDS-polyacrylamide gel electrophoresis (4-15% SDS-PAGE) and transferred onto methanol-activated polyvinylidene difluoride (PVDF) membrane (Bio-Rad). The following primary antibodies were used: anti-pIRF3 (S386) (1:1000) (Abcam), anti-total IRF3 (1:1000) (Immuno-Biological), anti-TRIM6 N-terminus (1:1000) (Sigma), anti-actin (1:2000) (Abcam), anti-pSTAT1 (Y701) (1:1000) (Cell Signaling), anti-pSTAT1 (S708) (1:2000), anti-total STAT1 (1:1000) (BD Biosciences), anti-VAMP8 (Cell Signaling) (1:500), anti-IKKε (T501) (1:1000) (Novus Biologicals), anti-IKKε (S172) (1:1000), anti-total IKKε (1:1000) (Abcam), anti-pTBK1 (S172) (1:1000) (Epitomics), anti-total TBK1 (1:1000) (Novus Biologicals), anti-total Jak 1 (BD Biosciences), and anti-pJak1 (Y1034/1035) (Cell Signaling). Immunoblots were developed with the following secondary antibodies: ECL anti-rabbit IgG horseradish peroxidase conjugated whole antibody from donkey (1:10,000), and ECL anti-mouse IgG horseradish peroxidase conjugated whole antibody from sheep (1:10,000) (GE Healthcare; Buckinghamshire, England). The proteins were visualized with either Pierce^TM^ or SuperSignal^®^ West Femto Luminol chemiluminescence substrates (Thermo Scientific). The amount of protein expressed was determined using Fiji (58) to calculate the area under the curve.

### IFNβ ELISA

Irradiated supernatants from WNV infected wt or TRIM6-KO A549 cells were collected at 8, 24, and 48 hpi to measure IFNβ using the VeriKine^TM^ Human IFN Beta ELISA kit (pbl assay science) following the manufacturer’s instructions. Standards were used to generated a standard curve to extrapolate the amount of IFNβ (pg/mL) in the supernatants. The limit of detection for the assay is 50pg/mL.

### Transcription Factor Binding Site Analysis

To identify transcription factors with binding sites within the VAMP8 genetic region, the VAMP8 RefSeqGene (NCBI: NG_022887.1) was analyzed with PROMO (Version 3.0.2). The transcription factors predicted within a dissimilarity margin less than or equal to 15% were identified and aligned to the VAMP8 RefSeqGene. Transcription factors with confirmed binding sites within the VAMP8 RefSeqGene were searched with JASPAR (2018) to confirm the transcription factor consensus sequence matched the sequence within *Vamp8*.

### Statistical Analysis

All analyses were performed in Graphpad Prism. All experiments were performed in triplicate. Statistical tests and measures of statistical significance are specified in the relevant figure legends. Repeated measures two-way ANOVA with Bonferroni’s post-test was applied for kinetics two factor comparisons (kinetics experiments), one-way ANOVA with Tukey’s post-test was used for comparing three more groups, and a student’s t-test for comparing two groups.

## REFERENCES

1. Gubler DJ, Kuno G, Markoff L. 2007. Flaviviruses, p 1153-1252, Fields Virology, 5 ed. Lippincott Williams & Wilins Publishers, Philadelphia, PA.

2. Lindenbach BD, Thiel HJ, Rice CM. 2007. Flaviviridae: The Viruses and Their Replication. Lippincott Williams & Wilins Publishers, Philadelphia, PA.

3. Kramer LD, Styer LM, Ebel GD. 2008. A global perspective on the epidemiology of West Nile virus. Annu Rev Entomol 53:61–81.

4. Anonymous. 2019-06-11T07:50:59Z/ 2019. West Nile virus | West Nile Virus | CDC. https://www.cdc.gov/westnile/index.html. Accessed

5. Sejvar JJ. 2014. Clinical manifestations and outcomes of West Nile virus infection. Viruses 6:606–23.

6. Dayan GH, Pugachev K, Bevilacqua J, Lang J, Monath TP. 2013. Preclinical and clinical development of a YFV 17 D-based chimeric vaccine against West Nile virus. Viruses 5:3048–70.

7. Dayan GH, Bevilacqua J, Coleman D, Buldo A, Risi G. 2012. Phase II, dose ranging study of the safety and immunogenicity of single dose West Nile vaccine in healthy adults >/= 50 years of age. Vaccine 30:6656–64.

8. Tesh RB, Arroyo J, Travassos Da Rosa AP, Guzman H, Xiao SY, Monath TP. 2002. Efficacy of killed virus vaccine, live attenuated chimeric virus vaccine, and passive immunization for prevention of West Nile virus encephalitis in hamster model. Emerg Infect Dis 8:1392–7.

9. Blazquez AB, Vazquez-Calvo A, Martin-Acebes MA, Saiz JC. 2018. Pharmacological Inhibition of Protein Kinase C Reduces West Nile Virus Replication. Viruses 10.

10. Morrey JD, Taro BS, Siddharthan V, Wang H, Smee DF, Christensen AJ, Furuta Y. 2008. Efficacy of orally administered T-705 pyrazine analog on lethal West Nile virus infection in rodents. Antiviral Res 80:377–9.

11. Perwitasari O, Cho H, Diamond MS, Gale M, Jr. 2011. Inhibitor of kappaB kinase epsilon (IKK(epsilon)), STAT1, and IFIT2 proteins define novel innate immune effector pathway against West Nile virus infection. J Biol Chem 286:44412–23.

12. Gorman MJ, Poddar S, Farzan M, Diamond MS. 2016. The Interferon-Stimulated Gene Ifitm3 Restricts West Nile Virus Infection and Pathogenesis. J Virol 90:8212–25.

13. Mertens E, Kajaste-Rudnitski A, Torres S, Funk A, Frenkiel MP, Iteman I, Khromykh AA, Despres P. 2010. Viral determinants in the NS3 helicase and 2K peptide that promote West Nile virus resistance to antiviral action of 2’,5’-oligoadenylate synthetase 1b. Virology 399:176–85.

14. Lazear HM, Diamond MS. 2015. New insights into innate immune restriction of West Nile virus infection. Curr Opin Virol 11:1–6.

15. Daffis S, Samuel MA, Suthar MS, Gale M, Jr., Diamond MS. 2008. Toll-like receptor 3 has a protective role against West Nile virus infection. J Virol 82:10349–58.

16. Daffis S, Lazear HM, Liu WJ, Audsley M, Engle M, Khromykh AA, Diamond MS. 2011. The naturally attenuated Kunjin strain of West Nile virus shows enhanced sensitivity to the host type I interferon response. J Virol 85:5664–8.

17. Errett JS, Suthar MS, McMillan A, Diamond MS, Gale M, Jr. 2013. The essential, nonredundant roles of RIG-I and MDA5 in detecting and controlling West Nile virus infection. J Virol 87:11416–25.

18. Lazear HM, Pinto AK, Vogt MR, Gale M, Jr., Diamond MS. 2011. Beta interferon controls West Nile virus infection and pathogenesis in mice. J Virol 85:7186–94.

19. Larena M, Lobigs M. 2017. Partial dysfunction of STAT1 profoundly reduces host resistance to flaviviral infection. Virology 506:1–6.

20. Lim JK, Lisco A, McDermott DH, Huynh L, Ward JM, Johnson B, Johnson H, Pape J, Foster GA, Krysztof D, Follmann D, Stramer SL, Margolis LB, Murphy PM. 2009. Genetic variation in OAS1 is a risk factor for initial infection with West Nile virus in man. PLoS Pathog 5:e1000321.

21. Bigham AW, Buckingham KJ, Husain S, Emond MJ, Bofferding KM, Gildersleeve H, Rutherford A, Astakhova NM, Perelygin AA, Busch MP, Murray KO, Sejvar JJ, Green S, Kriesel J, Brinton MA, Bamshad M. 2011. Host genetic risk factors for West Nile virus infection and disease progression. PLoS One 6:e24745.

22. Zhang HL, Ye HQ, Liu SQ, Deng CL, Li XD, Shi PY, Zhang B. 2017. West Nile Virus NS1 Antagonizes Interferon Beta Production by Targeting RIG-I and MDA5. J Virol 91.

23. Lubick KJ, Robertson SJ, McNally KL, Freedman BA, Rasmussen AL, Taylor RT, Walts AD, Tsuruda S, Sakai M, Ishizuka M, Boer EF, Foster EC, Chiramel AI, Addison CB, Green R, Kastner DL, Katze MG, Holland SM, Forlino A, Freeman AF, Boehm M, Yoshii K, Best SM. 2015. Flavivirus Antagonism of Type I Interferon Signaling Reveals Prolidase as a Regulator of IFNAR1 Surface Expression. Cell Host Microbe 18:61–74.

24. Laurent-Rolle M, Boer EF, Lubick KJ, Wolfinbarger JB, Carmody AB, Rockx B, Liu W, Ashour J, Shupert WL, Holbrook MR, Barrett AD, Mason PW, Bloom ME, Garcia-Sastre A, Khromykh AA, Best SM. 2010. The NS5 protein of the virulent West Nile virus NY99 strain is a potent antagonist of type I interferon-mediated JAK-STAT signaling. J Virol 84:3503–15.

25. Schuessler A, Funk A, Lazear HM, Cooper DA, Torres S, Daffis S, Jha BK, Kumagai Y, Takeuchi O, Hertzog P, Silverman R, Akira S, Barton DJ, Diamond MS, Khromykh AA. 2012. West Nile virus noncoding subgenomic RNA contributes to viral evasion of the type I interferon-mediated antiviral response. J Virol 86:5708–18.

26. Keller BC, Fredericksen BL, Samuel MA, Mock RE, Mason PW, Diamond MS, Gale M, Jr. 2006. Resistance to alpha/beta interferon is a determinant of West Nile virus replication fitness and virulence. J Virol 80:9424–34.

27. Sharma S, tenOever BR, Grandvaux N, Zhou GP, Lin R, Hiscott J. 2003. Triggering the interferon antiviral response through an IKK-related pathway. Science 300:1148–51.

28. Paul A, Tang TH, Ng SK. 2018. Interferon Regulatory Factor 9 Structure and Regulation. Front Immunol 9.

29. Fu XY, Kessler DS, Veals SA, Levy DE, Darnell JE, Jr. 1990. ISGF3, the transcriptional activator induced by interferon alpha, consists of multiple interacting polypeptide chains. Proc Natl Acad Sci U S A 87:8555–9.

30. Tenoever BR, Ng SL, Chua MA, McWhirter SM, Garcia-Sastre A, Maniatis T. 2007. Multiple functions of the IKK-related kinase IKKepsilon in interferon-mediated antiviral immunity. Science 315:1274–8.

31. Rajsbaum R, Versteeg GA, Schmid S, Maestre AM, Belicha-Villanueva A, Martinez-Romero C, Patel JR, Morrison J, Pisanelli G, Miorin L, Laurent-Rolle M, Moulton HM, Stein DA, Fernandez-Sesma A, tenOever BR, Garcia-Sastre A. 2014. Unanchored K48-linked polyubiquitin synthesized by the E3-ubiquitin ligase TRIM6 stimulates the interferon-IKKepsilon kinase-mediated antiviral response. Immunity 40:880–95.

32. Antonin W, Holroyd C, Tikkanen R, Honing S, Jahn R. 2000. The R-SNARE endobrevin/VAMP-8 mediates homotypic fusion of early endosomes and late endosomes. Mol Biol Cell 11:3289–98.

33. Wang CC, Ng CP, Lu L, Atlashkin V, Zhang W, Seet LF, Hong W. 2004. A role of VAMP8/endobrevin in regulated exocytosis of pancreatic acinar cells. Dev Cell 7:359–71.

34. Wang CC, Shi H, Guo K, Ng CP, Li J, Gan BQ, Chien Liew H, Leinonen J, Rajaniemi H, Zhou ZH, Zeng Q, Hong W. 2007. VAMP8/endobrevin as a general vesicular SNARE for regulated exocytosis of the exocrine system. Mol Biol Cell 18:1056–63.

35. Behrendorff N, Dolai S, Hong W, Gaisano HY, Thorn P. 2011. Vesicle-associated membrane protein 8 (VAMP8) is a SNARE (soluble N-ethylmaleimide-sensitive factor attachment protein receptor) selectively required for sequential granule-to-granule fusion. J Biol Chem 286:29627–34.

36. Jones LC, Moussa L, Fulcher ML, Zhu Y, Hudson EJ, O’Neal WK, Randell SH, Lazarowski ER, Boucher RC, Kreda SM. 2012. VAMP8 is a vesicle SNARE that regulates mucin secretion in airway goblet cells. J Physiol 590:545–62.

37. Loo LS, Hwang LA, Ong YM, Tay HS, Wang CC, Hong W. 2009. A role for endobrevin/VAMP8 in CTL lytic granule exocytosis. Eur J Immunol 39:3520–8.

38. Kanwar N, Fayyazi A, Backofen B, Nitsche M, Dressel R, von Mollard GF. 2008. Thymic alterations in mice deficient for the SNARE protein VAMP8/endobrevin. Cell Tissue Res 334:227–42.

39. Wang CC, Ng CP, Shi H, Liew HC, Guo K, Zeng Q, Hong W. 2010. A role for VAMP8/endobrevin in surface deployment of the water channel aquaporin 2. Mol Cell Biol 30:333–43.

40. Matheoud D, Moradin N, Bellemare-Pelletier A, Shio MT, Hong WJ, Olivier M, Gagnon E, Desjardins M, Descoteaux A. 2013. Leishmania evades host immunity by inhibiting antigen cross-presentation through direct cleavage of the SNARE VAMP8. Cell Host Microbe 14:15–25.

41. Ng SL, Friedman BA, Schmid S, Gertz J, Myers RM, Tenoever BR, Maniatis T. 2011. IkappaB kinase epsilon (IKK(epsilon)) regulates the balance between type I and type II interferon responses. Proc Natl Acad Sci U S A 108:21170–5.

42. Jiang D, Weidner JM, Qing M, Pan XB, Guo H, Xu C, Zhang X, Birk A, Chang J, Shi PY, Block TM, Guo JT. 2010. Identification of five interferon-induced cellular proteins that inhibit west nile virus and dengue virus infections. J Virol 84:8332–41.

43. Danial-Farran N, Eghbaria S, Schwartz N, Kra-Oz Z, Bisharat N. 2015. Genetic variants associated with susceptibility of Ashkenazi Jews to West Nile virus infection. Epidemiol Infect 143:857–63.

44. Zhang LK, Chai F, Li HY, Xiao G, Guo L. 2013. Identification of host proteins involved in Japanese encephalitis virus infection by quantitative proteomics analysis. J Proteome Res 12:2666–78.

45. Bharaj P, Atkins C, Luthra P, Giraldo MI, Dawes BE, Miorin L, Johnson JR, Krogan NJ, Basler CF, Freiberg AN, Rajsbaum R. 2017. The Host E3-Ubiquitin Ligase TRIM6 Ubiquitinates the Ebola Virus VP35 Protein and Promotes Virus Replication. J Virol 91.

46. Bharaj P, Wang YE, Dawes BE, Yun TE, Park A, Yen B, Basler CF, Freiberg AN, Lee B, Rajsbaum R. 2016. The Matrix Protein of Nipah Virus Targets the E3-Ubiquitin Ligase TRIM6 to Inhibit the IKKepsilon Kinase-Mediated Type-I IFN Antiviral Response. PLoS Pathog 12:e1005880.

47. Uchil PD, Hinz A, Siegel S, Coenen-Stass A, Pertel T, Luban J, Mothes W. 2012. TRIM protein-mediated regulation of inflammatory and innate immune signaling and its association with antiretroviral activity. J Virol 87:257–72.

48. van Tol S, Hage A, Giraldo MI, Bharaj P, Rajsbaum R. 2017. The TRIMendous Role of TRIMs in Virus-Host Interactions. Vaccines (Basel) 5.

49. Versteeg GA, Rajsbaum R, Sanchez-Aparicio MT, Maestre AM, Valdiviezo J, Shi M, Inn KS, Fernandez-Sesma A, Jung J, Garcia-Sastre A. 2013. The E3-ligase TRIM family of proteins regulates signaling pathways triggered by innate immune pattern-recognition receptors. Immunity 38:384–98.

50. Versteeg GA, Benke S, Garcia-Sastre A, Rajsbaum R. 2014. InTRIMsic immunity: Positive and negative regulation of immune signaling by tripartite motif proteins. Cytokine Growth Factor Rev 25:563–76.

51. Schmid S, Mordstein M, Kochs G, Garcia-Sastre A, Tenoever BR. 2010. Transcription factor redundancy ensures induction of the antiviral state. J Biol Chem 285:42013–22.

52. Daffis S, Szretter KJ, Schriewer J, Li J, Youn S, Errett J, Lin TY, Schneller S, Zust R, Dong H, Thiel V, Sen GC, Fensterl V, Klimstra WB, Pierson TC, Buller RM, Gale M, Jr., Shi PY, Diamond MS. 2010. 2’-O methylation of the viral mRNA cap evades host restriction by IFIT family members. Nature 468:452–6.

53. Olofsson P, Nerstedt A, Hultqvist M, Nilsson EC, Andersson S, Bergelin A, Holmdahl R. 2007. Arthritis suppression by NADPH activation operates through an interferon-beta pathway. BMC Biol 5:19.

54. Quiroga AD, Alvarez Mde L, Parody JP, Ronco MT, Frances DE, Pisani GB, Carnovale CE, Carrillo MC. 2007. Involvement of reactive oxygen species on the apoptotic mechanism induced by IFN-alpha2b in rat preneoplastic liver. Biochem Pharmacol 73:1776–85.

55. Fink K, Martin L, Mukawera E, Chartier S, De Deken X, Brochiero E, Miot F, Grandvaux N. 2013. IFNbeta/TNFalpha synergism induces a non-canonical STAT2/IRF9-dependent pathway triggering a novel DUOX2 NADPH oxidase-mediated airway antiviral response. Cell Res 23:673–90.

56. Dobin A, Davis CA, Schlesinger F, Drenkow J, Zaleski C, Jha S, Batut P, Chaisson M, Gingeras TR. 2013. STAR: ultrafast universal RNA-seq aligner. Bioinformatics 29:15–21.

57. Love MI, Huber W, Anders S. 2014. Moderated estimation of fold change and dispersion for RNA-seq data with DESeq2. Genome Biol 15:550.

58. A. Schindelin J, Arganda-Carreras I, Frise E, Kaynig V, Longair M, Pietzsch T, Preibisch S, Rueden C, Saalfeld S, Schmid B, Tinevez JY, White DJ, Hartenstein V, Eliceiri K, Tomancak P, Cardona 2012. Fiji: an open-source platform for biological-image analysis. Nat Methods 9:676–82.

